# Fluctuation spectroscopy of giant unilamellar vesicles using confocal and phase contrast microscopy

**DOI:** 10.1101/2020.05.20.104240

**Authors:** Hammad A. Faizi, Cody J. Reeves, Vasil N. Georgiev, Petia M. Vlahovska, Rumiana Dimova

## Abstract

A widely used method to measure the bending rigidity of bilayer membranes is fluctuation spectroscopy, which analyses the thermally-driven membrane undulations of giant unilamellar vesicles recorded with either phase-contrast or confocal microscopy. Here, we analyze the fluctuations of the same vesicle using both techniques and obtain consistent values for the bending modulus. We discuss the factors that may lead to discrepancies.

---

Bending rigidity of cellular membranes plays a key role in membrane remodeling. Knowledge of its value is needed to quantify processes that involve curvature changes such as budding (as in endo- and exocytosis), tubulation and fusion. Various experimental methods have been devised to measure bending rigidity[1], e.g. micropipette aspiration [2, 3], electrodeformation [5–8], optical tweezers [9, 10], and scattering based techniques [11, 12]. One of the most popular methods is fluctuation spectroscopy, pioneered by *Brochard and Lenon*[13], due to its ease of implementation[7, 14–16]. In essence, a time series of vesicle contours in the focal plane (the equator of the quasi-spherical vesicle) is recorded. The quasi-circular contour is decomposed in Fourier modes. The fluctuating amplitudes have variance dependent on the membrane bending rigidity and tension. Imaging is most commonly done by phase contrast microscopy [1, 7, 15, 17–22] but other methods such as confocal [23–25] and light sheet microscopy [26] have also been employed. The increased variety of imaging methods raises the question whether they all yield the same results.

Recently, *Rautu et al*. [25] pointed out that in phase contrast imaging, projections of out-of-focus fluctuations may contribute to the contour statistics leading to systematic overestimation of the bending rigidity value when compared to other methods such as micropipette aspiration and X-ray scattering. However, comparing bending rigidity numbers obtained by different techniques is only meaningful if the same system is probed. It is known that many factors such as sugars (and gravity), salt, buffers, solution asymmetry, concentration of fluorescent lipids, preparation method or type of bilayer configuration (stacked or free-floating), influence the measured mechanical properties of bilayer membranes [1, 18, 27–31]; see Table 1 in the Supporting Information (SI) for a list of reported bending rigidity of DOPC membranes. For example, even measurements with the same method can give a wide range of values, e.g., the bending rigidity of a DOPC bilayer measured with flickering spectroscopy has been reported from 15 k_*B*_*T*[24] to 30 k_*B*_*T* [20], where k_*B*_*T* is the thermal energy.

In order to compare imaging with phase contrast and confocal microscopy, which was suggested in *Rautu et al*. [25] as a better technique due to the precise control over the focal depth, we measure the bending rigidity of the same giant vesicle with both techniques. We highlight some important issues to be considered to ensure reliable measurements. We also show that results obtained with both methods are consistent.

## Equilibrium fluctuations of a quasi—spherical vesicle

First, we summarize the theoretical basis of the fluctuations analysis (details are provided in SI section S6). We also correct published expressions for the relaxation frequency and cross-spectral density of the shape fluctuations.

The contour in the equatorial plane of a quasi-spherical vesicle is decomposed in Fourier modes, 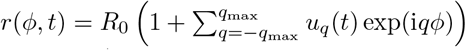, where *R*_0_ = (3*V*/4*π*)^1/3^ is the radius of an equivalent sphere with the volume *V* of the GUV and *q* is the mode number. In practice, *q*_max_ is the maximum number of experimentally resolved modes. The statistical analysis of the fluctuating amplitudes *u_q_* yields the values of membrane bending rigidity *κ* and the tension *σ* since 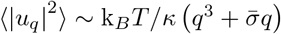, where 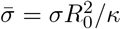.

More precisely, the statistics of the two-dimensional circular modes, *u_q_*, is derived from the three-dimensional shape modes, *f_lm_*, which describe the nearly-spherical shape in terms of spherical harmonics [14, 33], 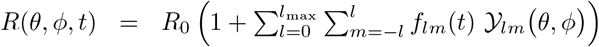. Here, *l*_max_ is an upper cutoff, in the order of the ratio of the GUV radius and bilayer thickness. The contour in the focal plane corresponds to the equator of the quasi-spherical vesicle, *θ* = *π*/2, i.e., 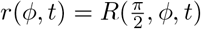, which leads to the following expression for the mean squared amplitudes

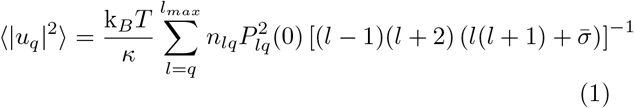

where *n_lq_* = (2*l* + 1)(*l* − *q*)!/4*π*(*l* + *q*)! and *P_lq_* are the associated Legendre polynomials. The short-wavelength shape fluctuations are dominated by the bending rigidity, while the long wavelengths are controlled by tension; the crossover occurs around mode 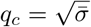.

To validate our methodology, we have simulated the thermal shape fluctuations of a GUV, see also SI section S6. We have generated a sequence of three-dimensional shapes (and their corresponding equatorial contours) using the evolution equations [33, 34]

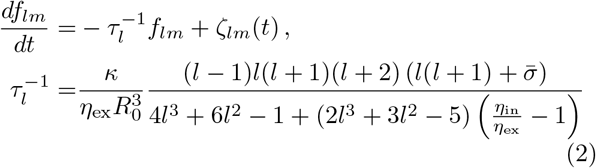

where *ζ_lm_* is the thermal noise driving the membrane undulations, *η*_in_ and *η*_ex_ are the viscosity of the solution inside and outside the vesicle. Note that the relaxation time given by Eq. 2 in *Rautu et al*. [25] has incorrect dependence on the viscosities of the enclosed and suspending solutions (this mistake is unlikely to affect the reported fluctuation spectra). The simulated contours were analyzed by our code and the extracted bending rigidity and tension were compared to the input values to confirm accuracy of the contour detection, Fourier decomposition and data fitting algorithms. The time evolution of the modes also enables us to access information provided by the time correlations 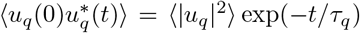. If *q* ≫ 1, the correlation time tends to that of a planar membrane 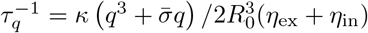. If the cross-spectral density 〈|*u_q_*(0)||*u_q_*(*t*)|〉 is utilized, the correct time dependence in the exponential includes a factor of 2 and Eq. 3 in *Zhou et al*. [35] needs to be corrected (see SI section S6, Eqn. 39).

## Bending rigidity obtained from confocal and phase-contrast microcopy: effect of resolution and vesicle size

Giant unilamellar vesicles (GUV) were electroformed from DOPC (99.8 mol% dioleoylphosphatidylcholine and 0.2 mol% Texas-Red 1,2-hexadecanoyl-wn-glycero-3-phosphoethanolamine, TR-DHPE) in 20 mM sucrose and subsequently diluted in 22 mM glucose, see SI section S2 for details. Low sugar concentration was used in order to minimize the effects of gravity[29] and effect of sugars[27], but still allow the vesicles to settle to the chamber bottom for easier recording. Low dye content minimizes effects of fluorophores[30].

Figure 1 shows a typical fluctuations spectrum, given by Eq. (1), fitted to the experimental data for the same vesicle imaged with confocal and phase contrast microscopy using a 40x objective with 0.6 numerical aperture (NA), pinhole size of 1 Airy unit (AU) and polarization correction (see below and SI section S3). The contour was detected with sub-pixel resolution [7]. The experimental data was fitted with Eq. (1) with Levenberg-Marquardt algorithm and yielded bending rigidity *κ* = 23.9 ± 1.6 k_*B*_*T* and tension *σ* = 5.1 ± 1.4 × 10^−9^ N/m and *κ* = 22.3 ± 2.1 k_*B*_*T* and *σ* = 3.1 ± 1.2 × 10^−9^ N/m from the confocal and phase contrast microscopy data, respectively. The average and error in individual GUV was determined by performing fluctuation spectroscopy 2-3 times. By imaging a population of 18 vesicles with both methods, the bending rigidity obtained are 22.5±2.0 k_*B*_*T* with confocal and 23.3±1.6 k_*B*_*T* phase contrast microscopy; each vesicle was analyzed with both imaging techniques as in Fig. 1 and then the results were averaged over the population. Figure 2 shows the box and whisker plot for more detailed statistics. Based on F statistics and ANOVA (analysis of variance) test, the *p*-value obtained is *p* = 0.48 for null hypothesis testing. We also performed the paired-sample t-test and obtained *p* = 0.43. These *p*-values indicate no significant difference between the two imaging techniques.

**FIG. 1.**
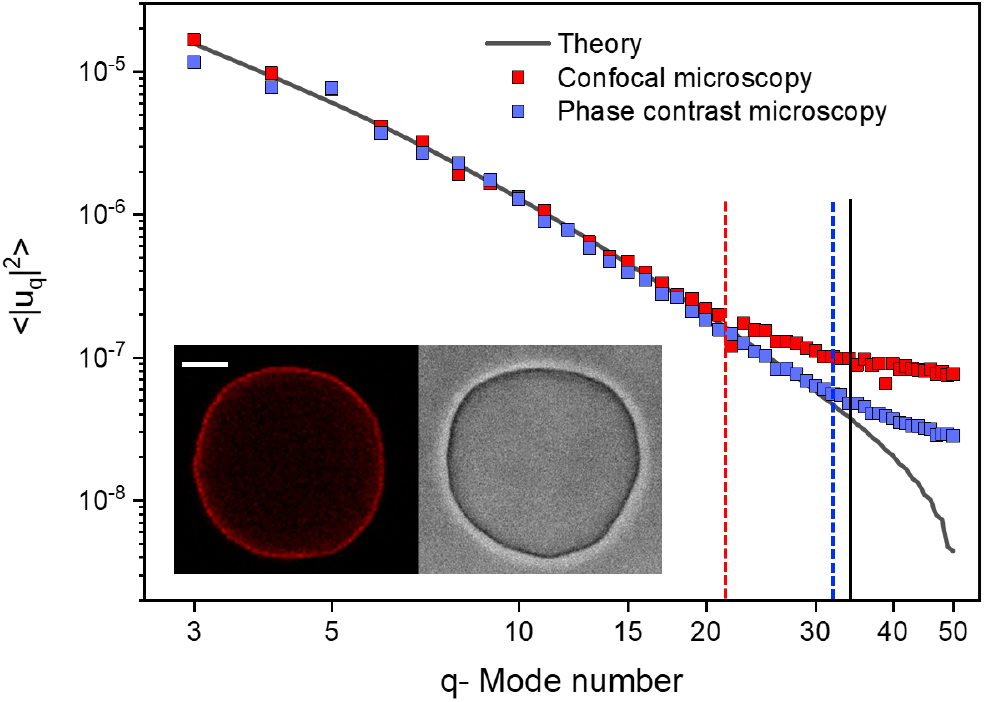
Fluctuation spectrum of the same DOPC vesicle (shown in the inset) obtained with confocal and phase contrast microscopy with a 40x/NA 0.6 objective, pinhole size of 1 AU and polarization correction, see also supplementary Movies S1 and S2. The dye concentration is 0.2 mol%. Scale bar is 15 *μ*m. The vertical lines denote the cutoff resolution for the modes: optical resolution (solid line), phase contrast (blue dashed) and confocal (red dashed line). The theory is fitted to phase contrast data up to the mode marked by the dashed blue line. The crossover mode 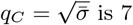 is 7.

**FIG. 2.**
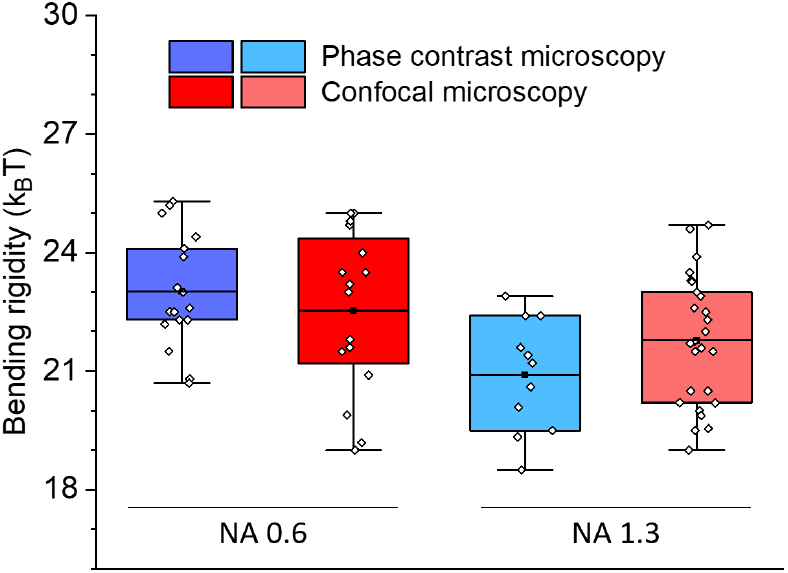
Imaging with phase contrast and confocal microscopy for objectives of the same numerical aperture (NA) give consistent results. Box and whisker plot comparison for a DOPC vesicle population where each vesicle was analyzed with phase contrast and confocal imaging with 40x objectives with NA 0.6 and NA 1.3. Pinhole size is 1 AU with polarization correction. The dye concentration is 0.2 mol%.

Since only modes with wavenumber 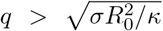 are sensitive to the bending rigidity, it is desirable to have more resolved modes, i.e., modes with amplitude and wavelength greater than the optical resolution limit ≈ 250 nm [36]. The average mean fluctuation amplitude scales with the size of the vesicle 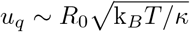, hence larger GUVs admit more spatially resolvable fluctuation modes as shown in Fig. 3. However, even for the same vesicle we find that the number of resolved modes is higher for phase contrast than for confocal imaging. Indeed, Fig. 1 shows that the noise level is higher for confocal microscopy, and on average phase contrast imaging resolves 8-10 modes more than confocal imaging does. The poorer mode resolution with confocal microscopy is likely due to poor contour recognition. The reasons for this are discussed in the next section.

**FIG. 3.**
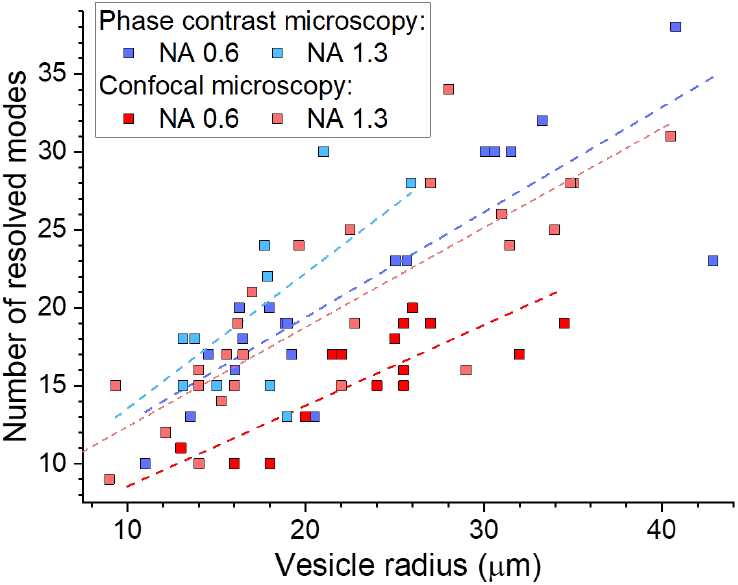
Larger vesicles allow resolving more fluctuation modes thus yielding more reliable determination of the bending rigidity. Data are collected on DOPC vesicles with different sizes. The dye concentration is 0.2 mol%. Regression lines are added to guide the eyes. Imaging was done with 40x objectives with different numerical aperture (NA), pinhole size of 1 AU and polarization correction.

We found that the vesicle population needs to have broad range of radii to avoid a size bias in the bending rigidity values we discovered for confocal microscopy with low resolution optics, see also SI section S4. In the case of 40x/NA 0.6 (air) objective, the mean Pearson correlation and standard deviation coefficient is 0.65±0.21 (see SI section S5 for the histograms generated with bootstrapping resampling technique). Analysis of a population of similar sized vesicles with radii around 10 *μ*m underestimates *κ* by roughly 6 k_*B*_*T*. The bias originates from out-of-plane fluorescence which worsens the contour detection. This issue is investigated in the next section and SI section 6. The size dependence is insignificant for phase contrast microscopy with a mean correlation coefficient of 0.28±0.18 with 40x/NA 0.6 (air). Analyzing the same vesicle population with 40x/NA 1.3 objective in phase contrast and confocal imaging yields 21.0±2.0 k_*B*_*T* and 21.7±2.0 k_*B*_*T* respectively. Higher numerical aperture in phase contrast leads to negligible correlation coefficient of −0.07±0.34 between bending rigidity and vesicle size and decrease in the correlation coefficient to 0.43±0.14 for confocal imaging with 40x/NA 1.3 objective.

## Out-of-focus fluorescence affects contour detection quality in confocal microscopy

The vesicle contour is detected from radial intensity line profiles, see SI section S2. In confocal cross sections, weak fluorescence from the vesicle membrane located above and below the focal plane may result in signal projected in the interior of the vesicle image which is higher compared to the surrounding background. The resulting asymmetry in the intensity line profile (Fig. 4a) leads to an artificial contour displacement, i.e., poor contour detection (note that such an asymmetry is absent in images acquired with phase contrast microscopy of vesicles with similar refractive indices of the internal and external solutions). This asymmetry creates a systematic error shifting the vesicle contour by 0.53 *μ*m. The error is larger than the pixel resolution of the system, 0.252 *μ*m, hence the higher modes are averaged out. Smaller vesicles or larger pinholes lead to higher signal inside the vesicle (see inset in Fig. 4b) corresponding to greater asymmetry which increases the error from contour fitting and introduces dependence of the bending rigidity on vesicle size. For imaging with higher numerical aperture objectives (e.g. NA 1.3), the asymmetry in the intensity line profiles is suppressed and contour detection is correct. Note that phase contrast images do not suffer from the asymmetry-induced error irrespective of the objective NA.

**FIG. 4.**
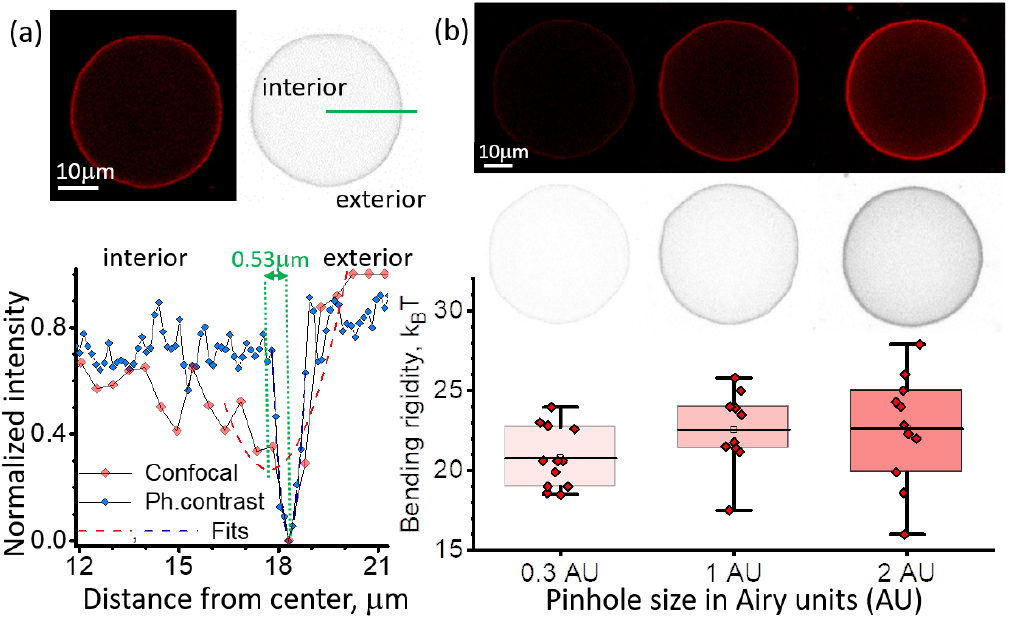
Out-of-focus fluorescence in confocal images can result in erroneous contour detection and increased error in bending rigidity. (a) Intensity line profiles (gray-value) across the vesicle membrane (DOPC, with 0.2 mol% dye) are symmetric for phase-contrast images (blue) but asymmetric for confocal images (red, 1 AU). The asymmetry in the confocal line profile leads to incorrect detection of the contour position defined by the parabolic fit minimum, here, shifted inwards by 0.53 *μ*m. (b) Vesicle images (and their inverted gray-value analogs) acquired with different pinhole size show increased fluorescence inside the vesicle which results in larger error in the bending rigidity. Box and whisker plot of the bending rigidity of the same DOPC vesicles imaged with confocal microscopy at three different pinhole sizes for 40x/NA 0.6 objective and polarization correction.

We investigated the impact of out-of-focus fluorescence on the fluctuations statistics by varying the pinhole size for confocal imaging on the same vesicle. The standard pinhole size in confocal microscopy is defaulted to 1 Airy unit (AU) (full width at half maximum FWHM=1.6 *μ*m) for 40x/NA 0.6 objective. We analyzed the same vesicles with different optical sectioning at 0.3 AU (FWHM=0.9 *μ*m), and 2 AU (FWHM=2.9 *μ*m). The mean bending rigidity did not show significant differences based on ANOVA testing, post hoc Dunnett test and paired-sample t-test (p=0.87), however the error increases with the pinhole size. The sensitivity to the vesicle size also becomes more pronounced with higher pinhole size. At the largest pinhole size (2.0 AU) the Pearson correlation coefficient is 0.60±0.22, while for 0.3 AU it becomes negligible, −0.14±0.30.

## Dye related artifacts: vesicle tubulation and polarization

Confocal imaging relies on fluorophores added to the membrane, and some studies have used up to 10 mol% dye [23, 24]. To probe the effect of fluorophore on *κ*, we changed the dye concentration from 0.2 mol% to 2 mol% TR-DHPE. The bending rigidity of this population of vesicles showed non significant difference with *κ*=20.09±2.49 k_*B*_*T* with one ANOVA testing. However, it was observed that over 2-3 min of recording, around 50 % of the vesicles developed inward structures such as buds or visible tubes as shown in Fig. 5. Vesicles with such defects displayed significantly higher bending rigidity, 25.01 ± 2.11 k_*B*_*T*.

**FIG. 5.**
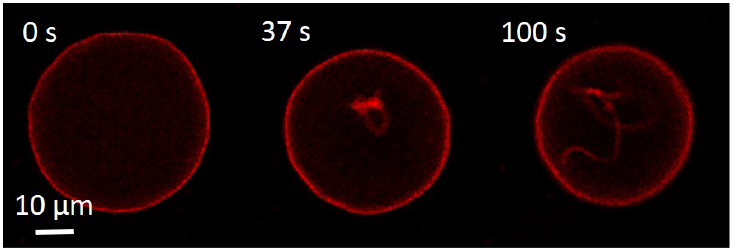
Time lapse of a DOPC vesicle with 2 mol% TR-DHPE developing inward nanotubes as a result of long exposure to laser during confocal imaging. The second and third cross sections are non-equatorial to better show the formed nan-otubes.

TR-DHPE belongs to a family of polarity-sensitive fluorescent probes. As a result, the signal intensities are different at the pole and equator of the vesicle (see SI section S3). This may lead to errors in the contour detection in these regions. The polarization effect was corrected by using circular rotation plates to have even intensities across the equatorial vesicle plane. The analysis of the same vesicle with and without the polarization correction showed a 3 k_*B*_*T* lower bending rigidity without any correction with 40x/NA 0.6 (air) objective. This softening effect became insignificant with 40x/NA 1.3 (oil) objective (SI section S3). This is likely due to loss of signal at low intensity regions where the higher mode fluctuations intensities are averaged out with background noise due to out-of-focus fluorescence.

## Effect of nearby vesicles on fluctuation spectra

The equilibrium shape fluctuations of an isolated GUV are driven by Gaussian thermal noise. Defects such as buds, nanotubes, invaginations or docked LUVs modify the vesicle fluctuations [7] and their effect can be detected in the statistics at each point on the vesicle contour profile using the ensemble-averaged probability density function (PDF) as shown in Fig. 6a In addition to defects attached the membrane, we also found that hydrodynamic flows and/or fluorescence signal from nearby vesicles can affect vesicle fluctuations.

**FIG. 6.**
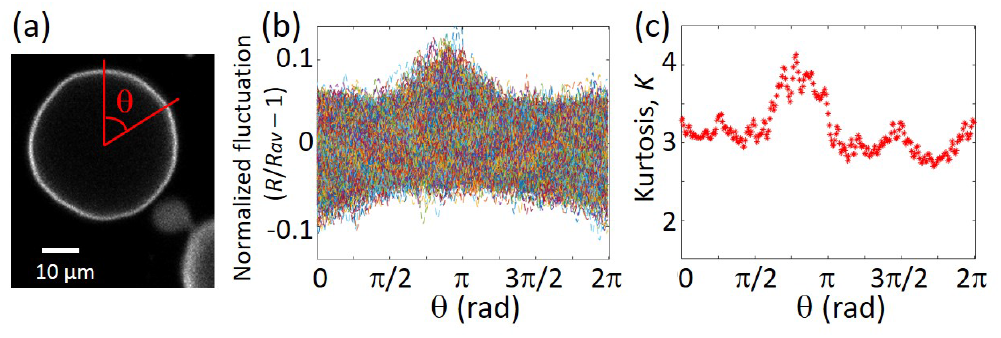
Nearby structures affect the fluctuation spectrum. (a) A flickering vesicle in close proximity to another vesicle bud. (b) Fluctuation density map of the vesicle in (a): the fluctuations are modified by hydrodynamic interactions of the other flickering vesicle bud. (c) Kurtosis *K* > 3 indicates the vesicle fluctuations have amplified meaning local softening of the membrane.

We characterized the Gaussianity of the fluctuations using the fourth PDF moment, Kurtosis, *K*. For a Gaussian distribution, *K* = 3. In Fig. 6 we demonstrate how thermal fluctuations may be modified (see supplementary Movie S3). As shown, the majority of contours are characterized by a normal distribution. However near other flickering structures, the fluctuation map density is modified. The non-Gaussian enhanced fluctuations are observed with leptokurtic nature (*K* > 3). This observation serves as a caution to filter out vesicles with sub-optical structures affecting the fluctuations.

## Conclusions

We compare the bending rigidity of bilayer membranes determined from flickering spectroscopy of GUVs imaged with confocal and phase contrast microscopy. Examining the same vesicle with both imaging techniques shows no significant differences in the bending modulus obtained from the two methods, in contrast to the overestimation reported by Rautu et al [25] when phase contrast microscopy is used. Our analysis indicates that membrane defects such as buds and tubes induced by long laser exposure in confocal microscopy can significantly stiffen the membrane. Furthermore, we find that errors in contour detection that could impact data interpretation can arise from fluorescence signal “pollution” and dye polarization. The bending rigidity we obtain (~ 22k_*B*_*T* for DOPC) is in line with the values obtained with other techniques such as micropipette aspiration, X-ray scattering, electrodeformation and neutron spin echo [3, 9, 11]. A scatter of approximately 2 k_*B*_*T* is typical in the experiments and should be taken into account when comparing data from different groups and methods. Exploring the effect of various parameters, we find that optimal imaging conditions for bending rigidity measurements from confocal imaging include high magnification objective, high numerical aperture, circular polarization correction, minimum dye concentration, small pinhole size, and broad vesicle size distribution.

In conclusion, we demonstrate that phase contrast and confocal microscopy produce the same results if precautions are taken to minimize effects of the dye and improve contour detection. Our study suggests that the many published results obtained by phase contrast microscopy are likely to be unaffected by the projections of out-of-focus fluctuations onto the imaging plane in contrast to the claim by Rautu et al [25]. Since dye related artifacts such as laser-induced defects can compromise the data, it is advantageous to use phase contrast imaging as it does not require dyes.

## ACKNOWLEDGEMENTS

This research was funded in part by NSF-CMMI awards 1748049 and 1740011. PV acknowledges support from the Alexander von Humboldt Foundation and HAF acknowledges financial support from Prof. Reinhard Lipowsky for visits to the Max Planck Institute of Colloids and Interfaces. We thank Paul Salipante for proofreading the manuscript.

## S1. BENDING RIGIDITY VALUE OF DOPC BILAYERS

The bending rigidity values of bilayer membranes made of the same lipid can vary across studies due to different conditions, e.g., sugars, salt, buffers, dye concentration, as well as the preparation method [37]. Table I illustrates the wide range of reported values of the bending rigidity values DOPC bilayers. Refer to Table II for the bending rigidity values obtained in this study for different microscopy setting.

**TABLE I.** Different bending rigidity values for DOPC under different conditions and methods.PC, C and EP refer to phase contrast, confocal and epi-flourescent microscopies used respectively in Fluctuation spectroscopy.

| Method | Rigidity ($k_B T$) | Dye conc. (%mol) | Buffer, Sugar (inside/outside) | Salt | Preparation |
| --- | --- | --- | --- | --- | --- |
| Fluctuation Spec. [EP,C] | 14.9±0.4 [24] | 15.8 NBD PC | 100 mM Sucrose/100 mM Sucrose | N/A | Electroformation |
| Fluctuation Spec. [PC] | 26.4±2.4 [38] | 0 or 0.1 diIC18 | 10 mM Sucrose/10 mM Glucose | 0.1 mM NaCl | Electroformation |
| Fluctuation Spec. [PC] | 26.8±2.4 [20] | 1.0 Liss Rhod PE | 450 mM Sucrose/500 mM Glucose | N/A | Electroformation |
| Fluctuation Spec. [PC] | 29.8±2.4 [20] | 1.0 Liss Rhod PE | 450 mM Sucrose/500 mM Glucose | N/A | Phase Transfer |
| Fluctuation Spec. [EP] | 22.3±0.5 [39] | 0.12 Liss Rhod PE | 100 mM Sucrose/200 mM Sucrose | N/A | Electroformation |
| Fluctuation Spec. [PC] | 27.3±3.2 [40] | N/A | 100 mM Sucrose/100 mM Sucrose | 2 mM NaN<sub>3</sub> | Electroformation |
| Fluctuation Spec. [PC] | 22.7±2 [41] | N/A | 100 mM Sucrose/125 mM Glucose | N/A | Electroformation |
| Fluctuation Spec. [PC] | 21.46±4 [42] | N/A | 100 mM Sucrose/125 mM Glucose | N/A | Electroformation |
| Fluctuation Spec. [PC] | 19±1 [43] | N/A | 10 mM Sucrose/10 mM Glucose | N/A | Electroformation |
| Fluctuation Spec. [C] | 19±1 [44] | 0.8 TR DHPE | 197 mM Sucrose/200 mM Glucose | N/A | Electroformation |
| Time Correlations | 22.1 [35] | N/A | 300 mM Sucrose/307 mM Glucose | N/A | Electroformation |
| Micropipette Aspiration | 20.7±2 [45] | N/A | 100 mM Sucrose/100 mM Glucose | N/A | Thin Film Hyd. |
| Micropipette Aspiration | 22.8±2.2 [43] | N/A | 8 mM Sucrose/8 mM Glucose | N/A | Electroformation |
| X- Ray Scattering | 20±2 [46] | N/A | Water/Water | N/A | Extrusion |
| X- Ray Scattering | 20.2±1.4 [47] | N/A | Deionized water | N/A | Bilayer stack |
| Electrodeformation | 21.9±2 [38] | 0.1 diIC18 | 10 mM Sucrose/10 mM Glucose | 0.1 mM NaCl | Electroformation |
| Tether pulling | 20±2 [10] | 0.3 TR DHPE | 300 mM Sucrose/80 mM Glucose | 100 mM NaCl | Electroformation |
| Neutron Spin Echo | 20±1 [48] | N/A | D<sub>2</sub>O/D<sub>2</sub>O | 0 mM | Extrusion |
| Neutron Spin Echo | 20±2 [49] | N/A | D<sub>2</sub>O/D<sub>2</sub>O | 0 mM | Extrusion |
| Neutron Spin Echo | 30±4 [49] | N/A | D<sub>2</sub>O/D<sub>2</sub>O | 150 mM | Extrusion |
| Neutron Spin Echo | 40±5 [49] | N/A | D<sub>2</sub>O/D<sub>2</sub>O | 470 mM | Extrusion |
| Interferometry | 10.5±8.8 [50] | N/A | 295 mM Sucrose/300 mM Glucose | N/A | Electroformation |

**TABLE II.**
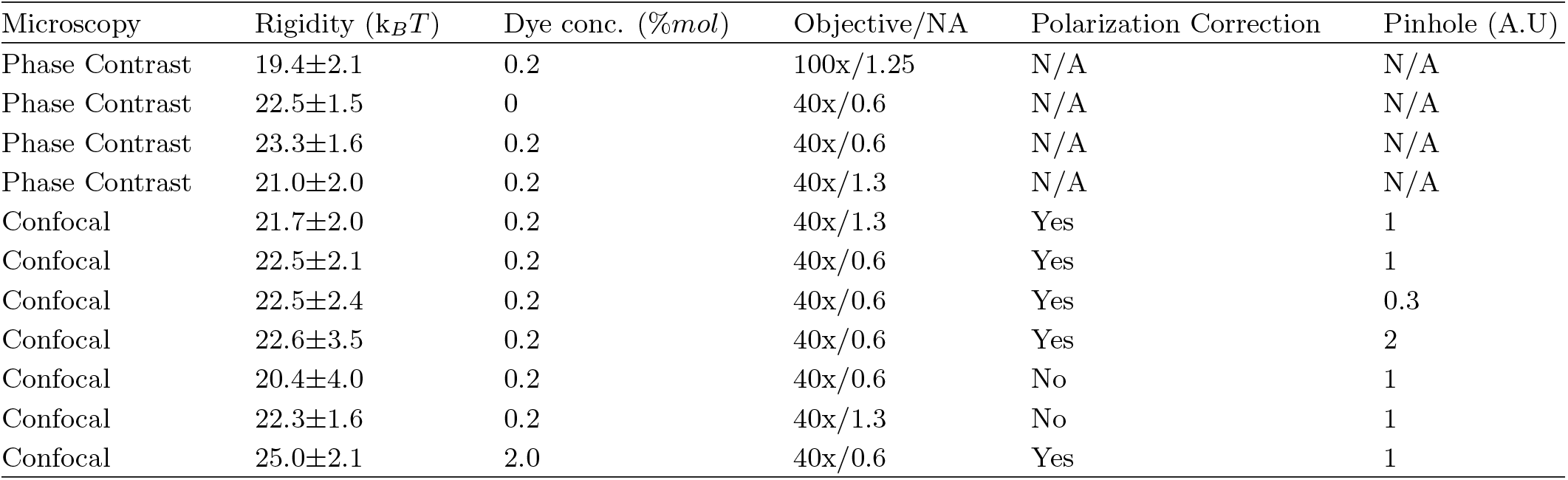
Bending rigidity values obtained in this study for DOPC under different conditions and microscopy settings. Note the sugar concentration is the same in all the experiments: 20 mM Sucrose inside/ 22 mM Glucose outside. The dye used is TR DHPE and all the vesicles were formed via electroformation.

## S2. METHODS

### Vesicle preparation

Giant unilamellar vesicles (GUVs) were prepared using the classical electroformation method [51] from DOPC and the fluorescent lipid Texas Red 1,2-hexadecanoyl-sn-glycero-3-phosphoethanolamine (TR-DHPE). The composition of the GUVs explored are 99.8 % DOPC 0.2 % TR-DHPE and 98 % DOPC 2 % TR-DHPE (mole fractions). Stock solutions of DOPC and TR-DHPE at 10 mg/ml and 1 mg/ml in chloroform were diluted to a final concentration of 4 mM for varying proportions. A small volume, 10 *μ*l, of the solution was spread on the conductive surface of two glass slides coated with indium tin oxide (ITO) (Delta Technologies). The glass slides were then stored under a vacuum for 1–2 hours to remove traces of organic solvent. Afterwards, a 2 mm Teflon spacer was sandwiched between the glass slides and the chamber was gently filled with 20 mM sucrose solution. The slides (conductive side facing inward) were connected to an AC signal generator Agilent 33220A (Agilent Technology GmbH, Germany). An AC field of voltage 1.5 V and frequency 10 Hz applied for 2 hours at room temperature, resulting in 10-50 *μ*m sized vesicles. The harvested vesicles were diluted 10 times in 22 mM glucose solution to obtain fluctuating vesicles. All GUVs were analyzed within 8 hours of electoformation.

### Microscopy and video recording

The equatorial fluctuations for both phase contrast and confocal mode were recorded with Leica TCS SP8 scanning confocal microscope using a HCX PL APO 40x/ Numerical Aperture (NA) 0.6 Ph2 (air) objective and a HC PL APO 40x/ NA 1.3 (oil) objective. The pinhole size during the experiment was fixed to 1 AU (Airy units) unless stated otherwise. Table 1 compiles the pixel size and focal depth for different experimental conditions. The scanning speed was fixed to 1 kHz in bidirectional mode and the polarizer plates were rotated (100%) to remove the polarization effect of the fluorescent dye unless stated otherwise. The dye was excited with a 561 nm laser (diode-pumped solid-state laser) with 1.61% (laser intensity) HyD3 detector (hybrid) and the gain was fixed to 23%. Phase contrast imaging was recorded with PCO CS dimax (PCO AG, Kelheim, Germany)) mounted on confocal microscope. 1500-2000 images were recorded at 3.83 frames per second (fps) with confocal and 60 fps with phase contrast imaging. The RGB confocal images were converted to 8 bit and then inverted. We implemented an inbuilt MATLAB sobel disk filter *f special*(‘*sobel*’) and image normalization to increase the contrast of the contour.

In this section, we list different focal depths and pixel sizes for different microscopy and numerical aperture settings for 40x objective. Focal depth or FWHM (full width half maximum) of phase contrast imaging was determined using the standard formula 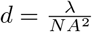 The wavelength of transmission light was assumed to be 550 nm.

**TABLE III.** Different experimental conditions for video recording with 40x objective.

| Microscopy | Numerical Aperture | Medium | Pinhole size (AU) | Focal depth ($\mu\text{m}$) | Pixel Resolution (nm) |
| --- | --- | --- | --- | --- | --- |
| Phase contrast | 0.6 | Air | 1 | 1.57 | 276.9 |
| Phase contrast | 1.3 | Oil | 1 | 0.35 | 158.7 |
| Confocal | 0.6 | Air | 1 | 1.61 | 252.7 |
| Confocal | 1.3 | Oil | 1 | 0.52 | 252.7 |

**TABLE IV.**
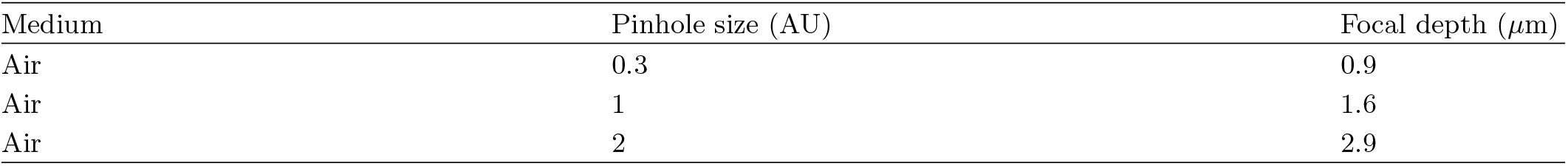
Focal depth or FWHM (full width half maximum) for confocal imaging.

| Medium | Pinhole size (AU) | Focal depth ($\mu\text{m}$) |
| --- | --- | --- |
| Air | 0.3 | 0.9 |
| Air | 1 | 1.6 |
| Air | 2 | 2.9 |

### Sub-pixel contour recognition

**FIG. 7.**
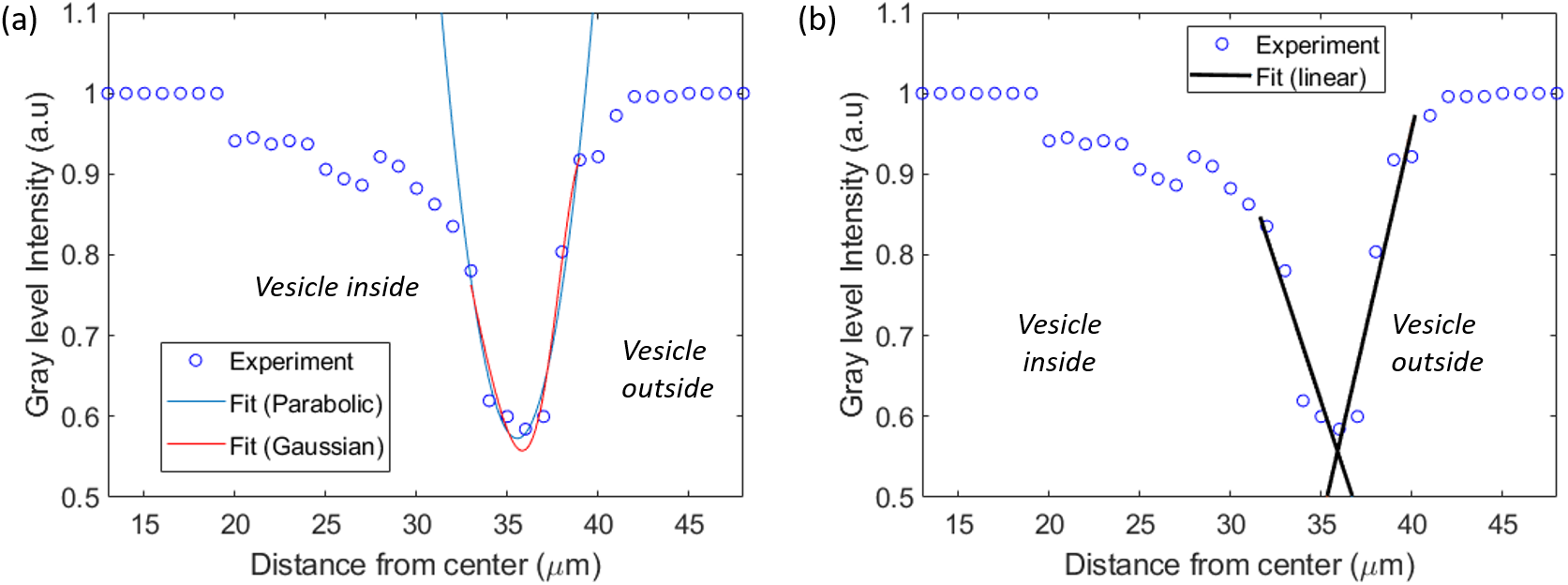
Intensity profile for a vesicle contour obtained from confocal imaging. The contour recognition details are given in [7]. The sub-pixel accuracy of the contour profile is determined based on (a) Gaussian, parabolic and (b) linear interpolations

The intensity profile in the radial direction for N wedges were determined from three different interpolation schemes (Gaussian, parabolic and linear weighting of neighbouring pixel) for sub-pixel contour recognition. This was done to check if different interpolation schemes affects the bending rigidity values due to uncertainty introduced at higher wave-numbers for experimental vesicle contour fluctuations. The mean bending rigidity obtained was similar for all the three schemes for the same vesicle. Figure (S7) illustrates the subpixel accuracy determination for a 35 *μ*m radius vesicle. The bending rigidities obtained was 22.0±3.0 k_*B*_*T*, 21.1±1.0 k_*B*_*T* and 21.9±2.2 k_*B*_*T* from Gaussian, parabolic and linear interpolation schemes respectively.

## S3. POLARIZATION EFFECTS

We analyzed the same vesicle with and without polarization effects. The polarization effects were corrected using circular plates that were rotated 100%. Figure (S8) illustrates the effect of dye polarization for vesicles imaged with different numerical apertures. Using one Anova test, we find a significant difference of 3 k_*B*_*T* for the 40x/0.6 NA case. The difference tends to be negligible for 40x/1.3 NA case.

**FIG. 8.**
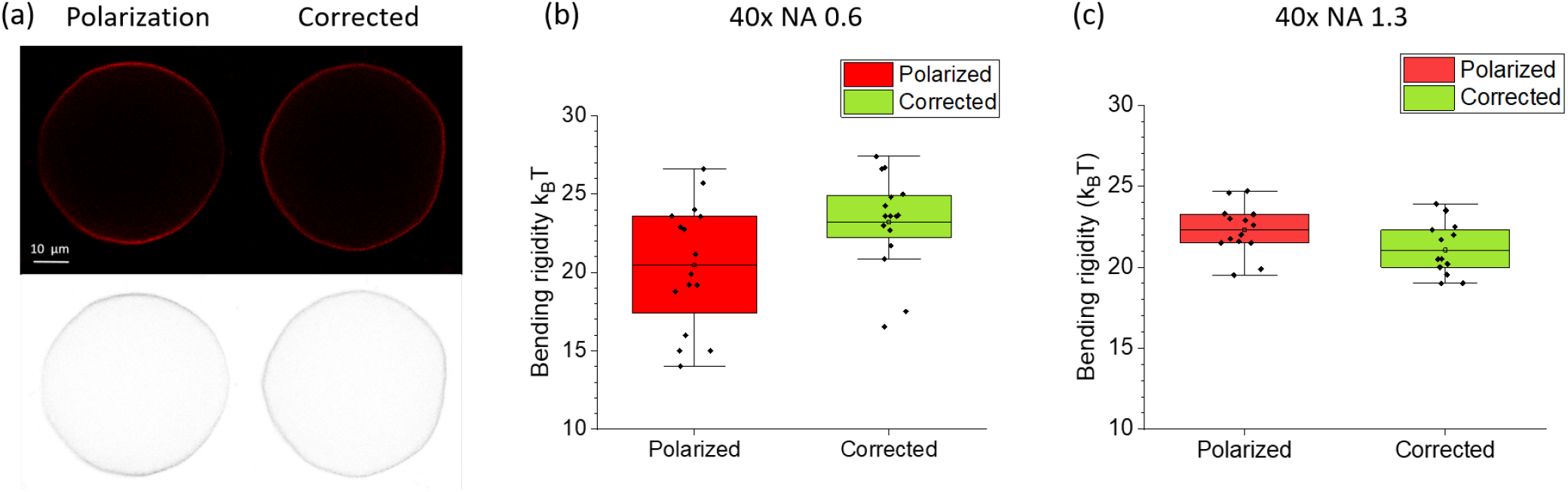
Polarization effects. (a) Confocal images of the same vesicle with and without polarization effects for 40x/0.6 NA case. The polarization effects were removed using circular plates that were rotated 100%. (b, c) Comparison between the same vesicles for different numerical apertures. Using one Anova test, we find a significant difference of 3 k_*B*_*T* for 40x/0.6 NA case. The difference tends to be negligible for 40x/1.3 NA case. Pinhole size is 1 AU.

## S4. EFFECT OF VESICLE SIZE ON BENDING RIGIDITY VALUES

The bending rigidity obtained from confocal microscopy with low-resolution optics (e.g. 40x objective, NA 0.6, 1 AU, polarization correction) can be systematically underestimated if the vesicle population contains similar sized vesicles. We demonstrate this by comparing the bending rigidity of the same vesicle imaged with confocal and phasecontrast microscopy, see Figure (S9). Vesicles with smaller sizes yield apparently lower bending rigidity, see Fig. S4 which further highlights the bias effect. For small vesicles, the out-of-focus signal gives rise to asymmetry in the contour intensity (illustrated in Fig. 4a in the main text) which leads to errors in the contour detection and underestimation of the bending rigidity. When the refractive index difference across the membrane is small (as is the case in our experiments), phase contrast imaging does not suffer from this size bias.

## S5. BOOTSTRAPPING RESAMPLING

Details about the various statistical techniques can be found in Ref. [52]. Here we explain the bootstrapping sampling technique. A more rigorous reference is the textbook [53]. In practice, the finite amount of data or length of experiment limits the accuracy to infer data confidently. Bootstrapping is an inference method about the population from a given sample. In bootstrap-resamples, the population is in fact the sample and this quantity is known. This allows to measure the quality of inference of the ‘true’ sample from a re-sampled data. For example, let’s consider the average mass of the human population world wide. It is difficult to measure the mass of every individual globally, therefore, a small sample is measured. Let’s assume the sample size of N people. From that sample size, only one mean can be measured. In order to have a reasonable estimate about the population statistics, we need to have variability of the mean that we computed. The simplest bootsampling statistics can be considered by taking the original data N individuals and resampling to create a new sample of the same size N (e.g. we might ‘resample’ 10 times from [60, 61, 62, 63, 64, 65, 66, 67] kg and get [61, 64, 63, 63, 60, 60, 62, 65] kg). This process is repeated a large number of times, 100 to 10000, to create a histogram that be applied to any estimator testing. Bootstrap resampling was carried out using MATLAB’s *bootstrp ()*.

**FIG. 9.**
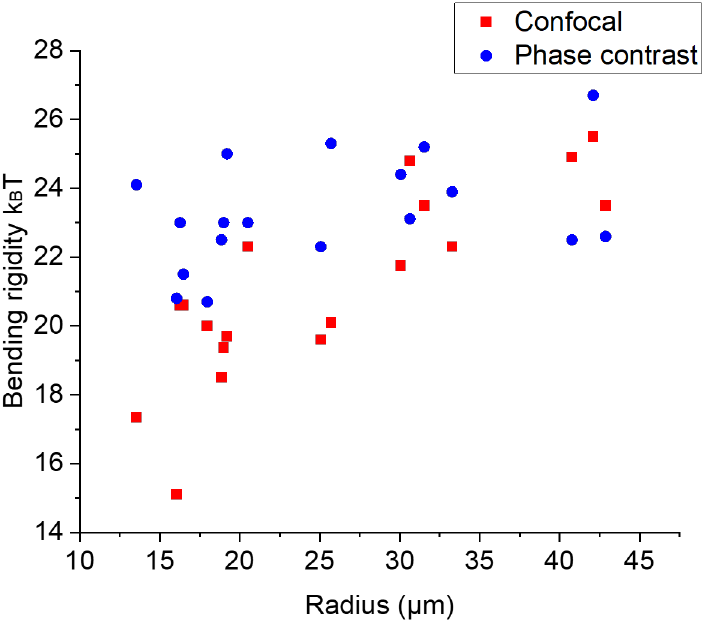
Vesicle size effects. Every vesicle was imaged with confocal and phase contrast microscopy. Data are collected on DOPC vesicles with different sizes. The dye concentration was 0.2 mol. Imaging was done with 40x objectives with NA 0.6, 1 AU and polarization correction.

**FIG. 10.**
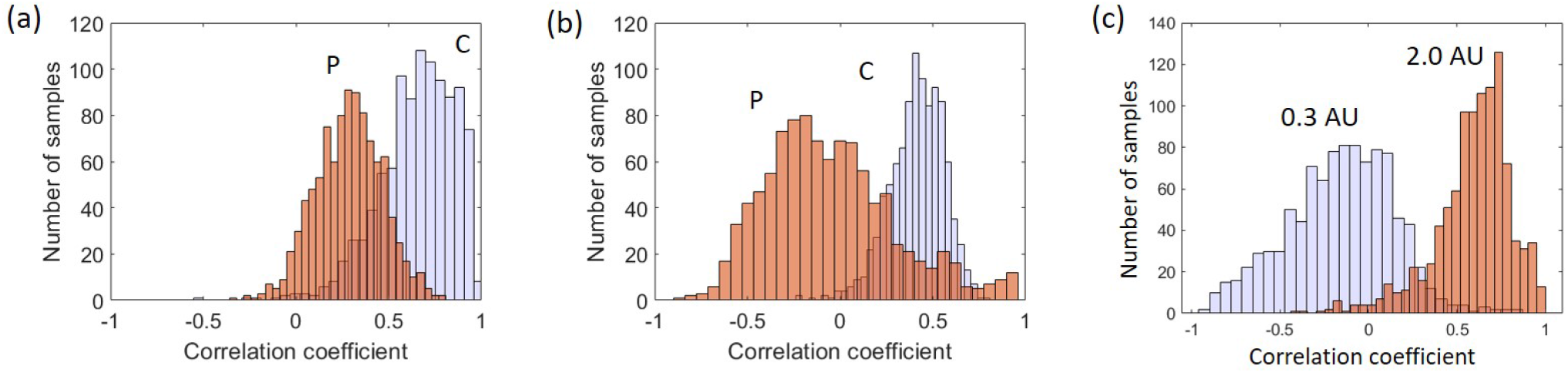
Bootstrap method with 95% confidence to evaluate bending rigidity dependence on size of vesicles for different numerical aperture (a) 40x/0.6 NA, (b) 40x/1.3 NA in phase contrast (P) or confocal(C) microcopy, and the pinhole sizes (c) (blue AU 0.3 and red AU 2).

In the case of our experiments, the finite amount of data or length of experiment limits the accuracy to infer data confidently. The bootstrap resampling requires choosing random replacement from a given data set and examining each sample the same way. This way a particular data point from the original set can reappear randomly multiple times in a particular bootstrap sample. The element size of the bootstrap sample is the same as the element size of the original data. This technique allows to obtain uncertainty of the quantity one estimates.

Bootstrap resampling algorithm for estimating standard error [53]:

1. Obtain N independent bootstrap samples *X*\*^1^, *X*\*^2^, *X*\*^3^,… *X*\*^*N*^, each consisting of n data values drawn with a replacement from *x* where *x* = [*x*^1^, *x*^2^, *x*^3^…*x^n^*]. Note for estimating a standard error, the number N will ordinarily be larger than 30 to satisfy the Central Limit Theorem. Computations allow to use a large number N such as 10^3^ to 10^4^.
2. Determine the bootstrap replication for every bootstrap resample:

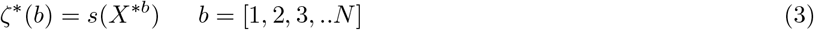

where *s*() is a statistical function like sample mean. For example, if *s*(*x*) is the sample mean 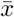 then *S*(*X*\*) is the mean of bootstrap data set.
3. Compute the standard error *SE* by utilizing the standard deviation of N replications

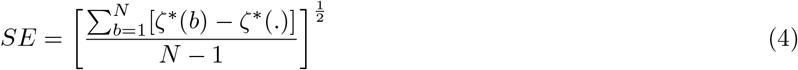

where 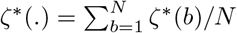.

In our case we determine the *SE* of mean Pearson correlation using bootsampling statistics.

## S6. NUMERICAL SIMULATIONS OF VESICLE CONTOURS

### Mathematical Model

The total energy of the system is given by the Helfrich model[54] as Eq. (5) where *κ* is the bending rigidity, *c*_1_ and *c*_2_ are the local radii curvatures, *A* is the total surface area, *V* is the interior volume of the vesicle, *σ* is the surface tension, and *p* is the pressure difference across the membrane.

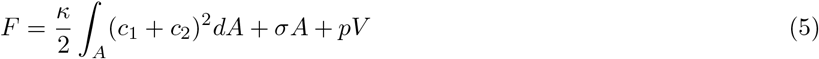

For a quasi-spherical vesicle in equilibrium, the shape can be decomposed into spherical harmonics 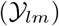 such that the position of the surface is given by

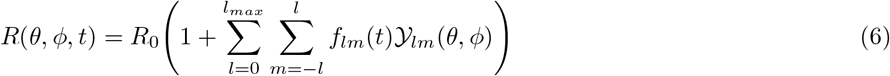

where the characteristic radius *R*_0_ is given by 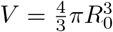. The spherical harmonics are defined as

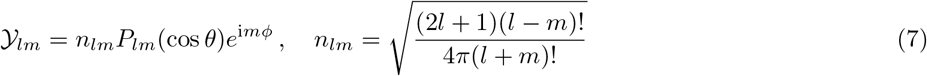

*P_lm_*(cos *θ*) are the associated Legendre polynomials.

As *l* = 1 account for translational modes, for the sake of this paper, *f_lm_*(*t*) will be restricted to *f*_1*m*_(*t*) = 0 for *l* = 1. Furthermore, volume conservation requires that [33]

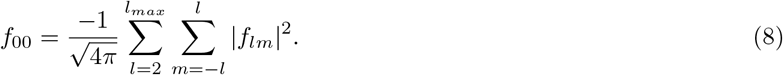

Assuming there is no external fluid flow, the harmonic coefficients (*f_lm_*) for *l* > 1 are described by the following

**FIG. 11.**
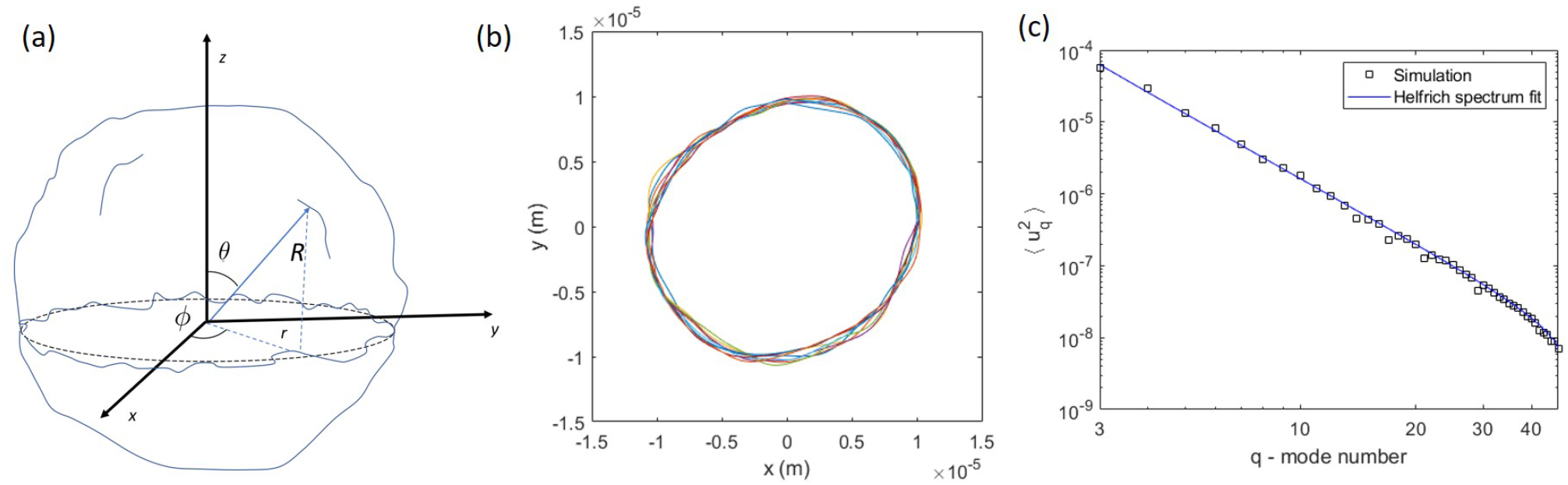
(a) A sketch of a GUV. (b) Time sequence of vesicle contours taken at time intervals of 1 s; the bending rigidity is *κ* =1× 10^−19^ J and the membrane tension is *σ* =1× 10^−9^ N/m. The size of the vesicle is *R*_0_ = 10^−5^ m. (c) Helfrich mode spectrum determined by the image detecting algorithm based on Ref. [7]. The spectrum was fitted with Equation 2 from the main text to obtain the bending rigidity and membrane tension.

stochastic differential equation [33]

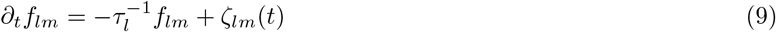

where

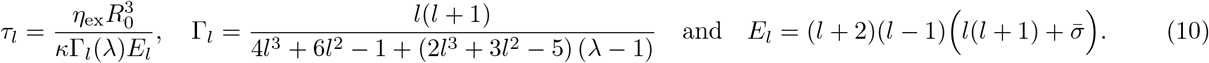

A tutorial derivation of the evolution equation and the relaxation time (in the absence of thermal noise) can be found in Refs. [56, 57] The dimensionless tension is 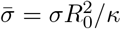. λ = *η*_in_/*η*_ex_ is the ratio of viscosities of the solutions inside and outside the vesicle. When λ =1, our result for the relaxation time reduces to the one reported by Refs. [33, 55]. To make easier comparison with the result of Ref. [44], we can rewrite the relaxation time as

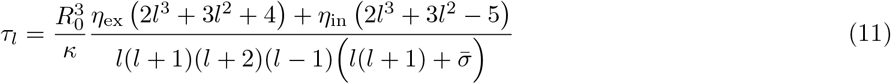

*ζ_lm_*(*t*) is a stochastic term accounting for thermal noise; the corresponding time correlation is given as

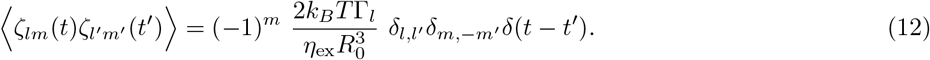

The *δ* functions are the traditional Kronecker and Dirac delta functions. From Eq. 12, the variance of *ζ_lm_*(*t*) is given by

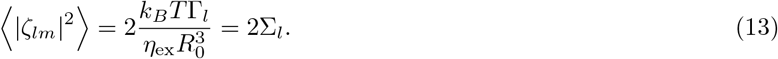

### Numerical Method

At this point, it is convenient to decompose *f_lm_* and *ζ_lm_* into real and imaginary components such that *f_lm_*(*t*) = *X_lm_*(*t*) + i *Y_lm_*(*t*) and *ζ_lm_*(*t*) = *a_lm_*(*t*) + i *b_lm_*(*t*). As *a_lm_* and *b_lm_* are independent of each other then

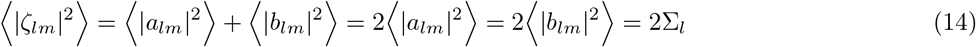

Eq. (9) can then be rewritten as

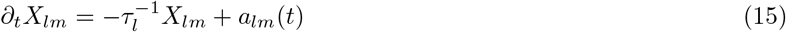

and similarly for *Y_lm_*. As Eq. (15) is a simple Langevin equation, the exact time update [?] is given as

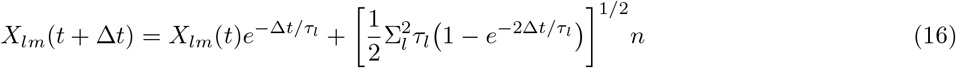

such that Δ*t* is the time step size and *n* is a sample value from the normal distribution 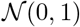. In order to properly resolve the dynamics of the higher order coefficient, a sufficiently small time step must be chosen so that Δ*t* << *τl*_max_. Yet as each harmonic coefficient is independent of each other, Eq. 16 can be evaluated for all *X_lm_* and *Y_lm_* simultaneously. Given all the harmonic coefficients (*f_lm_*), the cross-section at the equator, *R*(*θ* = *π*/2), can easily be computed using Eq. (6).

When running the numerical simulations, the user has some choice of which input parameters to specify. For example, one can specify the effective surface tension (dimensionless) 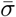 and the largest incorporated mode *l_max_*. In this case, the vesicle’s excess is obtained from [33]

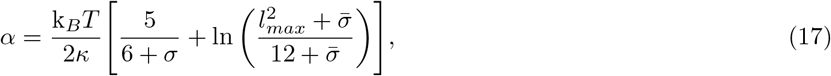

Alternatively, one can specify *α* and 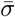, and Eq. (17) then provides the requisite *l_max_*.

Here we demonstrate an example of a numerically simulated vesicle with predefined bending rigidity and membrane tension. Figure (S11)b shows a time sequence of equatorial vesicle contours with bending rigidity of *κ* = 10^−19^ J and membrane tension of *σ* = 10^−9^ N/m. The size of the vesicle is *R*_0_ = 10^−5^ m. By implementing our image detection technique and fitting algorithm from *Gracia et al*. [7], we are able to reproduce the bending rigidity and membrane tension respectively as *κ* = (1.00 ± 0.01) × 10^−19^ J and *σ* = (1.1 ± 0.2) × 10^−9^ N/m with the Helfrich spectrum given in Figure (S11). Notably our image detection is able to resolve more than 45 shape fluctuation modes.

### Simulating the Effect of Out-of-focus Signal

Due to a finite focal depth, the microscope imaging does not capture only the optical/fluorescence signal at the focal (equatorial) plane. The out-of-focus signal results in gradient in the image intensity near the focal plane vesicle contour.

To simulate this effect, we numerically projected the vesicle shape *R*(*θ, ϕ, t*) on the equatorial plane and assigned intensity of the projected location, *R*(*θ, ϕ, t*) sin *θ*, given by

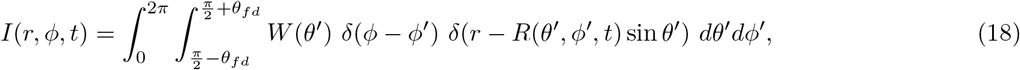

where 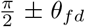 are the top and bottom of the microscope focal depth (*FD*), *θ_fd_* = arctan(*FD*/*R*_0_). *W*(*θ*) is the intensity weighting function

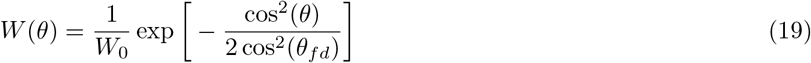

and *W*_0_ is the corresponding normalization constant. The resulting images of the equatorial plane at different focal depth are illustrated in Figure S12a.

We varied the magnitude of the focal depth *FD*, from 0 to 0.3*R*_0_. The fluctuation spectra obtained for the simulations are shown in Figure S12b for a vesicle sized *R*_0_ = 20*μm* with *κ*= 22 k_*B*_*T* and *σ* = 1.4×10^−9^ N/m. The crossover mode 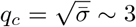. The effect of the projections is only significant for modes *q* ≥ Δ^−1^, where 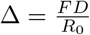 [44]. For smaller values of Δ < 0.05, the projections have no effect - the spectra overlap implying same bending rigidity. However, as the value of Δ increases, more modes get affected by the projections resulting in an effective softening of the membrane from 22 k_*B*_*T* to 19 k_*B*_*T*, see S12c.

### Fluctuations statistics: derivations of the basic results

Here we summarize the main results for the dynamics of a quasi-spherical vesicle.

#### Mean Squared Magnitude of the Fourier Modes

The dynamics of the spherical harmonics modes is governed by the following Langevin equation,

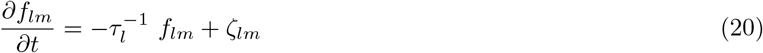

where the relaxation time *τ_l_* is given by Eq. (11) and the noise is

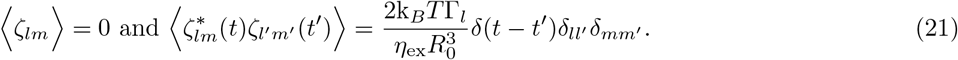

The analytic solution to Eq. (20) is given by

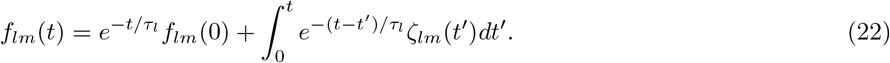

and therefore

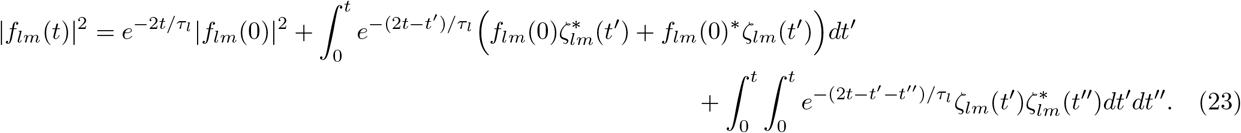

The ensemble average of 〈|*f_lm_*|^2^〉 of Eq. (23) is then

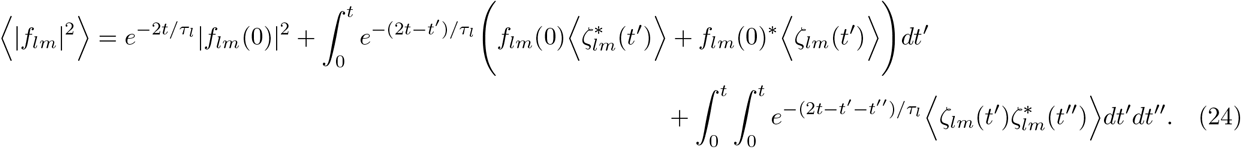

**FIG. 12.**
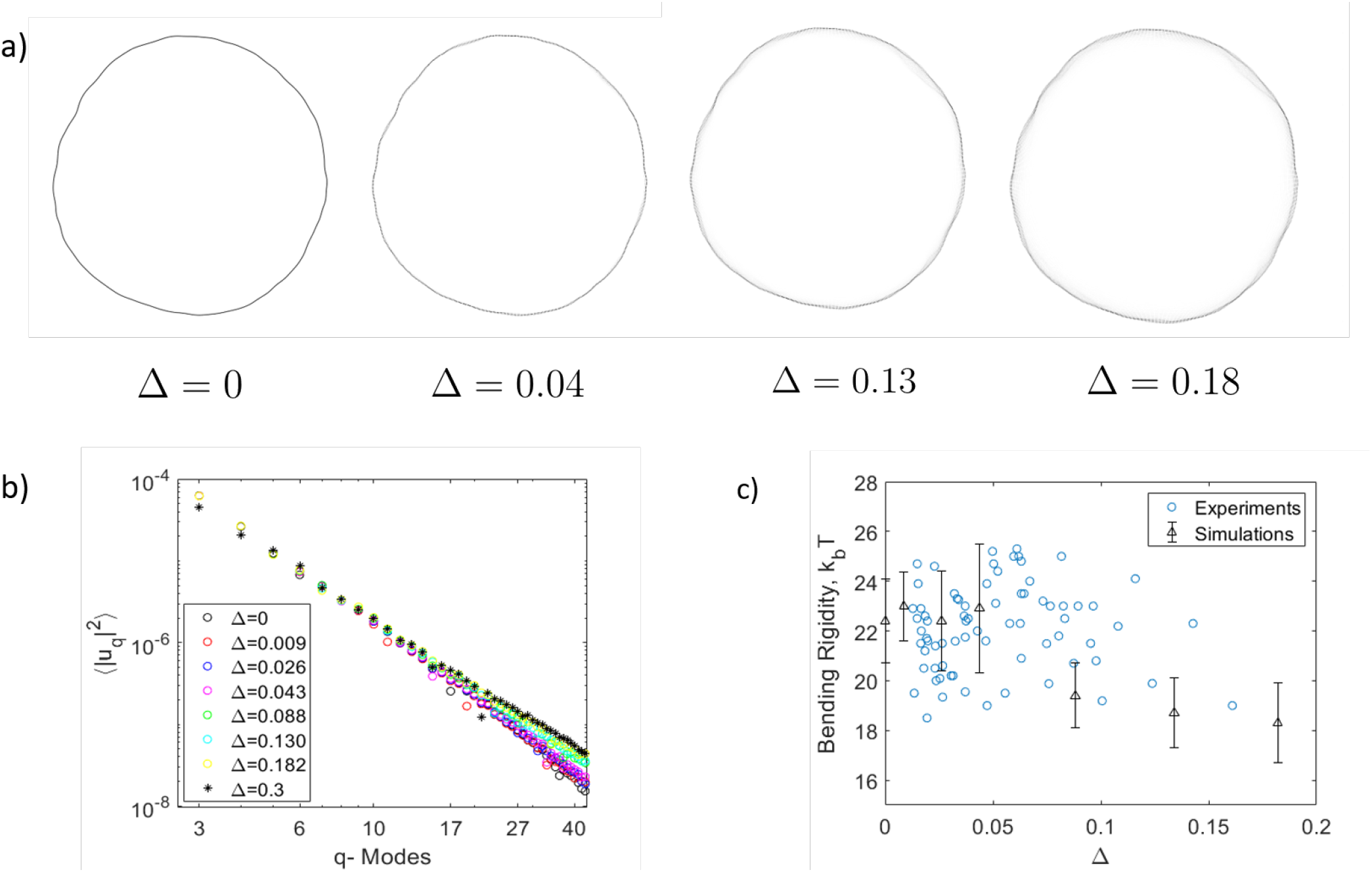
a) Snapshot of vesicle equatorial contours at different Δ = *FD*/*R*_0_. The simulated vesicle has bending rigidity= 22 k_*B*_*T*, membrane tension= 1.4×10^−9^ N/m and radius *R*_0_ = 20 *μm*. Each image was acquired over 0.2 s (corresponding to imaging rate of 5 fps). b) Fluctuation spectrum obtained at different Δ from the numerical simulations c) Bending rigidity obtained for different Δ. Here we have compared the experimental results with numerical simulations.

Using Eq. (21), Eq. (24) simplifies to

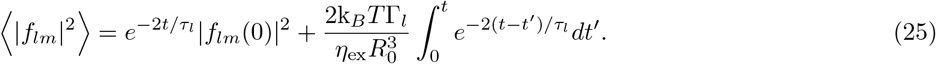

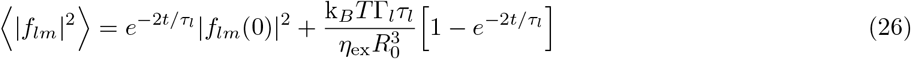

At long times, *t* >> *τ_l_*, Eq. (26) simplifies to

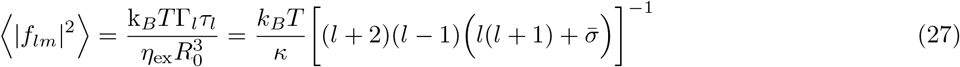

Recall 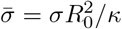. Since the dynamics of the different spherical harmonics modes are completely decoupled, we can more generally say

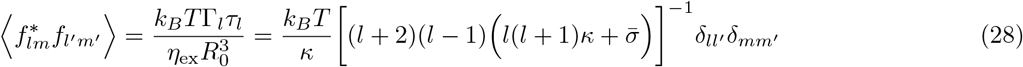

Next, we consider the contour of the GUV at the equator as a function of the spherical harmonic coefficients:

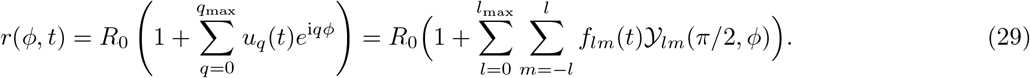

The Fourier coefficient for the *q*-th mode is then given by

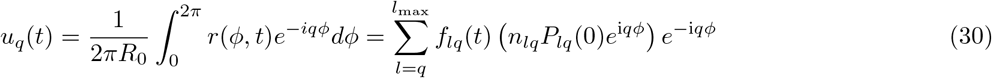

as all the other terms integrate to zero. In the above equation, we have inserted the definition of the spherical harmonic, 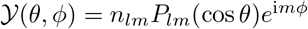 (see Eq. (7)), which shows that the dependence on *ϕ* cancels out.

The mean squared amplitude of *u_q_* is then given by

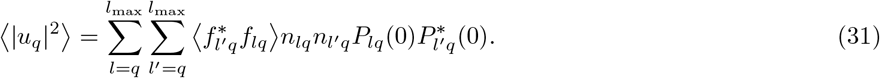

Using Eq. (28), the above equation simplifies to

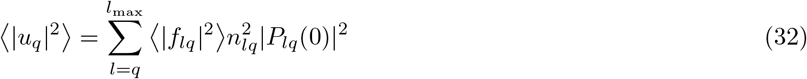

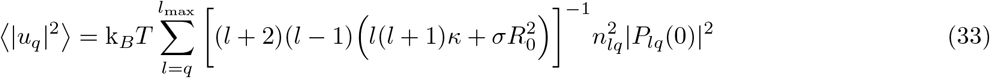

Eq. (33) follows *q*^−3^ behavior for bending dominated modes 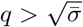 (and *q*^−1^ behavior for tension dominated modes 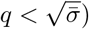).

#### Time Correlation for Fourier Modes

Time correlations present another useful metric to analyze the membrane fluctuations. As the different spherical harmonics modes are independent, the average time correlations,

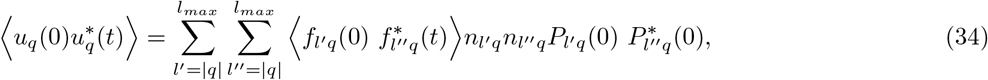

can be simplified to

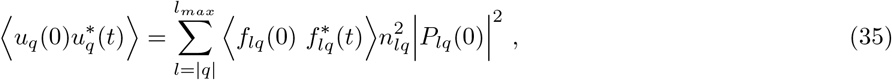

Using (23), (34) can be rewritten as

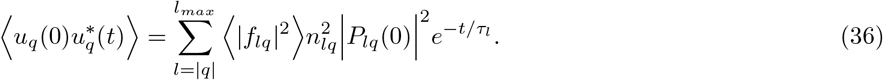

Since the first term in (36) has both the smallest decay rate 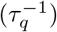 and largest mean-squared amplitude, the time correlation can be approximated to leading order as

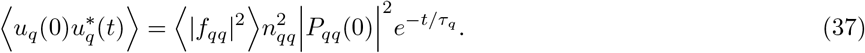

If we consider limit of undulations with short wavelengths (shorter than the vesicle radius), *q* ≫ 1, then the leading order decay rate can be approximated as

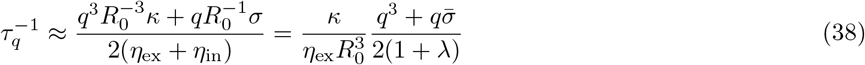

which is the decay rate derived using planar fluctuations. However, we suggest using the exact decay rate from the spherical harmonics as it is both more accurate and valid for all Fourier modes.

When comparing the time correlations in Fig. 13, the exact decay rate, from the full spherical harmonics (SpH), is immediately more accurate than if the planar membrane (PM) decay rate is used. To get the accuracy even better, the higher order terms in Eq. (36) must be included. If all of the terms are included then the time correlation is directly on top of the curve from produced by the numerical simulation. However, as it is not feasible to include all the terms for real membranes, it is of interest to know how many terms are enough to sufficiently reproduce the numerical simulations. As shown in Figure 5, the time correlation produced by including the first two terms in Eq. (36 lies almost directly on top of the true solution. Including more terms would improve the accuracy further, but it is not likely to be significant due to experimental error.

**FIG. 13.**
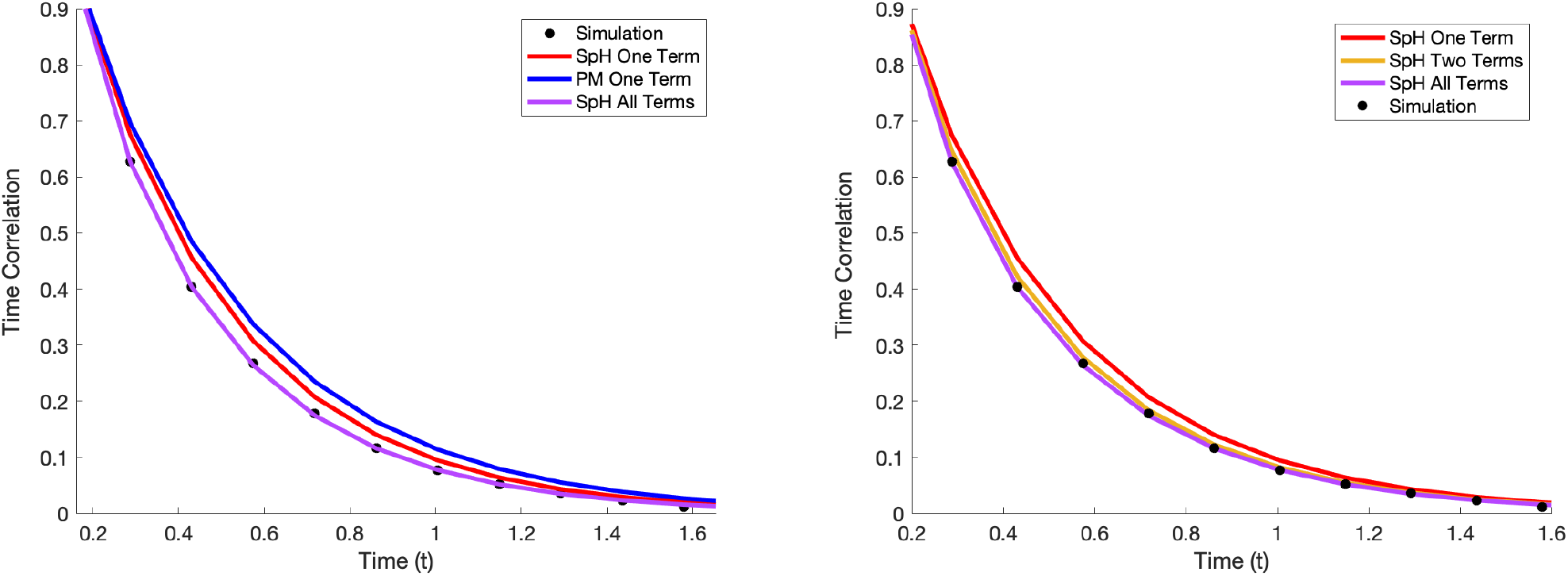
Plots comparing the analytic approximations for time correlation for Fourier mode *q* = 5. The left plots the time correlations using the exact spherical harmonic (SpH) decay rate and the less accurate planar membrane (PM) decay rate. The right plots the time correlations for the SpH case using different number of terms. The black dots show the time correlations computed from a numerical simulation using the following parameters as inputs. *R*_0_ =3 × 10^−5^ m, *κ* = 5 × 10^−19^ J, *σ* = 4 × 10^−8^ N/m, *l_max_* = 14

#### Cross-Spectral Density

Similar to time correlations, the Cross-Spectral Density (CSD) is given by

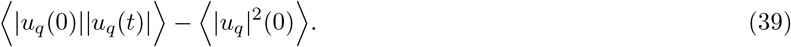

For the sake of clarity of explanation, in this section we will use the leading order approximation of *u_q_*,

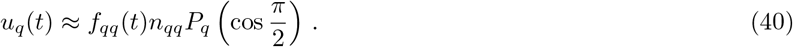

Using (22), this can be rewritten as

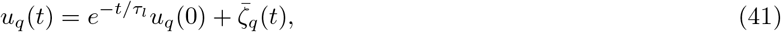

where

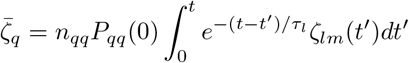

is a random normally distributed Weiner process.

From (41), it is clear that 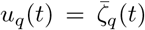 for large values of *t*. Furthermore, it is worth noting that all Fourier modes, except *q* = 0, have both real and an imaginary component, *u_q_* = *A_q_* + i*B_q_*, and that these two components are independent of each other. Likewise, the thermal noise can be decomposed into independent real and imaginary components: 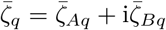. The real component of Eq. (41) can then be written as

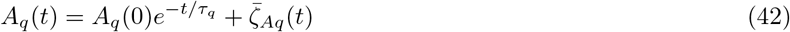

and a similar expression for *B_q_*.

Therefore, it can be shown that

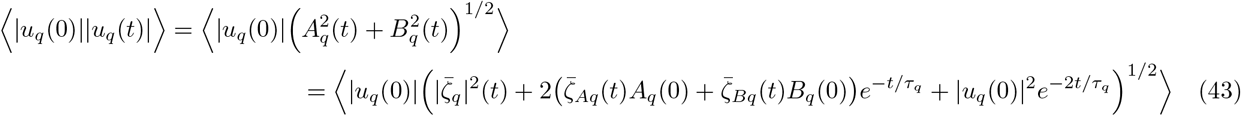

If we assume that *t* >> *t_q_*, then we can perform the following expansion

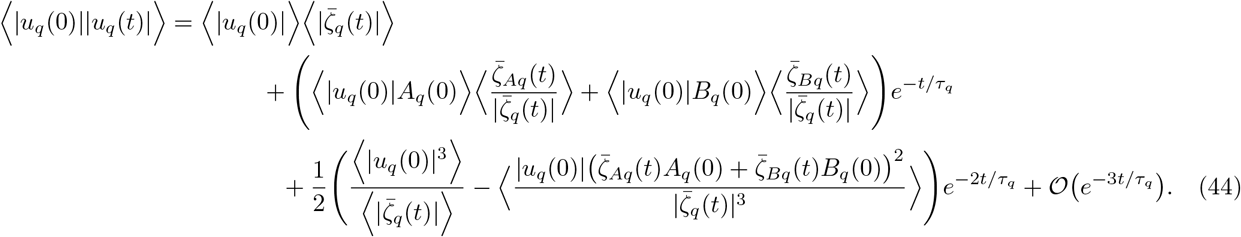

The second term in (44) averages to zero due to the thermal noise factor. Therefore, to leading order, the CSD is given as

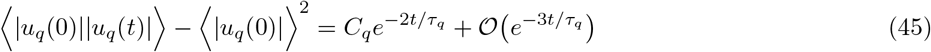

where *C_q_* is a normalization constant.

Therefore, the slowest decaying mode of the CSD is 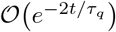. This contradicts H. Zhou et al. [35] who give it as 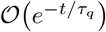. This factor of two is a consequence that each Fourier coefficient has both a real and imaginary component that are completely independent of each other.

Finally, users are recommended to use time correlations over CSD. CSD requires the same amount of work and contains the same higher order error as the time correlation method. Yet, CSD has an additional layer of truncation error introduced in the expansion in Eq. (44).

### Movie S1

Real-time video of the GUV from Figure 1 in the main text (DOPC labeled with 0.2 mol % TR-DHPE) acquired with phase contrast microscopy. The vesicle radius is 29.6 *μ*m.

### Movie 2

Video of the GUV from Figure 1 in the main text (DOPC labeled with 0.2 mol % TR-DHPE) acquired with confocal microscopy. The objective used is 40x/0.6 NA with the pinhole size 1 A.U. The polarization effect was corrected by using circular rotation plates to have even intensities across the equatorial vesicle plane. The vesicle radius is 29.6 *μ*m.

### Movie S3

Real time video of the GUV from Figure 6 in the main text consisting of DOPC labeled with TR DHPE (0.8%) acquired with confocal microscopy. The objective used is 40x/0.6 NA at 13.2 fps with the pinhole size 1 A.U. The polarization effect was corrected by using circular rotation plates to have even intensities across the equatorial vesicle plane.

### Raw data

Raw data are available at https://dx.doi.org/10.17617/3.4p. This collection of raw data, consists of 4 folders each containing zipped files of data.

In folder “Raw data - Fourier modes”, 2 different sets of experimental data are included: phase contrast (PC) and confocal (C) microscopy on the same vesicle and confocal microscopy data with different pinhole sizes. The folder contains excel sheets with fluctuation amplitude for every Fourier mode and the metadata with all the microscopy conditions. The vesicles have different sizes so that they practically cover a good span of focal depth (Δ) from 0.03 to 0.15. The meta data is included in the first sheet of the excel file. The second sheet has the mode and mean squared amplitude (and error). The remaining sheets have the Fourier modes for every microscopy setting (and focal depth). Note that our Fourier signal was normalized by vesicle radius. For the definition of our Fourier transform, please refer to *Gracia et al*. [7]. The contour detection is conducted as given in the main text and supplement.

The rest of the folders as listed below contain vesicle images in tiff format (grouped in folders for the separate vesicle as suggested by the folder name) and an excel sheet with the meta data indicating the specific microscopy conditions (AU = Airy unit, NA = numerical aperture).

The folder “Raw images - different focal depth” contains confocal microscopy raw images at different focal depths without any image processing for vesicles of different sizes.

The folder “Raw images - phase contrast vs confocal” contains phase contrast and confocal microscopy raw images for the same vesicle without any image processing for vesicles of different sizes.

The folder “Raw images - polarization correction” contains polarized and polarization-corrected confocal microscopy raw images for the same vesicle without any image processing.

## References

[1] Rumiana Dimova. Recent developments in the field of bending rigidity measurements on membranes. Advances in Colloid and Interface Science, 208:225–234, 2014. ISSN 0001-8686. doi: https://doi.org/10.1016/j.cis.2014.03.003. URL http://www.sciencedirect.com/science/article/pii/S0001868614001031. Special issue in honour of Wolfgang Helfrich.

[2] E. Evans and W. Rawicz. Entropy driven tension and bending elasticity in condensed-fluid membranes. Phys. Rev. Lett., 64:2094–2097, 1990.

[3] W. Rawicz, K.C. Olbrich, T. McIntosh, D. Needham, and E. Evans. Effect of chain length and unsaturation on elasticity of lipid bilayers. Biophysical Journal, 79(1):328–339, 2000. ISSN 0006-3495. doi:https://doi.org/10.1016/S0006-3495(00)76295-3. URL http://www.sciencedirect.com/science/article/pii/S0006349500762953.

[4] E. Parra-Ortiz and D. Needham. Mechanic assays of synthetic lipid membranes based on micropipette aspiration. In R. Dimova and C. Marques, editors, The Giant Vesicle Book, page Chapter 11. CRC Press, 2019.

[5] M. Kummrow and W. Helfrich. Deformation of giant lipid vesicles by electric fields. Phys. Rev. A, 44:8356–8360, 1991.

[6] P. M. Vlahovska, R. S. Gracia, S. Aranda-Espinoza, and R. Dimova. Electrohydrodynamic model of vesicle deformation in alternating electric fields. Biophys. J., 96: 4789–4803, 2009.

[7] Rubèn Serral Gracia, Natalya Bezlyepkina, Roland L. Knorr, Reinhard Lipowsky, and Rumiana Dimova. Effect of cholesterol on the rigidity of saturated and unsaturated membranes: fluctuation and electrodeformation analysis of giant vesicles. Soft Matter, 6:1472–1482, 2010. doi: 10.1039/B920629A. URL http://dx.doi.org/10.1039/B920629A.

[8] P. F. Salipante, R. Knorr, R. Dimova, and P. M. Vlahovska. Electrodeformation method for measuring the capacitance of bilayer membranes. Soft Matter, 8:3810–3816, 2012.

[9] Aiwei Tian, Benjamin R. Capraro, Cinzia Esposito, and Tobias Baumgart. Bending stiffness depends on curvature of ternary lipid mixture tubular membranes. Biophysical Journal, 97(6):1636–1646, 2009. ISSN 0006-3495. doi:https://doi.org/10.1016/j.bpj.2009.07.012. URL http://www.sciencedirect.com/science/article/pii/S0006349509012338.

[10] B. Sorre, A. Callan-Jones, J.-B. Manneville, P. Nassoy, J.-F. Joanny, J. Prost, B. Goud, and P. Bassereau. Curvature-driven lipid sorting needs proximity to a demixing point and is aided by proteins. PNAS, 106: 5622–5626, 2009.

[11] Norbert Kučerka, Yufeng Liu, Nanjun Chu, Horia I. Petrache, Stephanie Tristram-Nagle, and John F. Nagle. Structure of fully hydrated fluid phase dmpc and dlpc lipid bilayers using x-ray scattering from oriented multilamellar arrays and from unilamellar vesicles. Biophysical Journal, 88(4):2626–2637, 2005. ISSN 0006-3495. doi: https://doi.org/10.1529/biophysj.104.056606. URL http://www.sciencedirect.com/science/article/pii/S0006349505733178.

[12] Jianjun Pan, Frederick A. Heberle, Stephanie Tristram-Nagle, Michelle Szymanski, Mary Koepfinger, John Katsaras, and Norbert Ku?erka. Molecular structures of fluid phase phosphatidylglycerol bilayers as determined by small angle neutron and x-ray scattering. Biochimica et Biophysica Acta (BBA) - Biomembranes, 1818(9):2135–2148, 2012. ISSN 0005-2736. doi: https://doi.org/10.1016/j.bbamem.2012.05.007. URL http://www.sciencedirect.com/science/article/pii/S0005273612001551.

[13] Brochard, F. and Lennon, J.F. Frequency spectrum of the flicker phenomenon in erythrocytes. J. Phys. France, 36(11):1035–1047, 1975. doi: 10.1051/jphys:0197500360110103500. URL https://doi.org/10.1051/jphys:0197500360110103500.

[14] JF Faucon, MD Mitov, P Meleard, I Bivas, and P Bothorel. Bending Elasticity and Thermal Fluctuations of lipid membranes - Theoretical and Experimental requirements. Journal De Physique, 50(17): 2389–2414, SEP 1 1989. ISSN 0302-0738. doi: 10.1051/jphys:0198900500170238900.

[15] J. Pecreaux, H.-G. Dobereiner, J. Prost, J.-F. Joanny, and P. Bassereau. Refined contour analysis of giant unilamellar vesicles. Eur. Phys. J. E., 13:277–290, 2004.

[16] Julia Genova, Victoria Vitkova, and Isak Bivas. Registration and analysis of the shape fluctuations of nearly spherical lipid vesicles. Phys. Rev. E, 88:022707, Aug 2013. doi:10.1103/PhysRevE.88.022707. URL https://link.aps.org/doi/10.1103/PhysRevE.88.022707.

[17] Patricia Bassereau, Benoit Sorre, and Aurore Levy. Bending lipid membranes: Experiments after w. helfrich’s model. Advances in Colloid and Interface Science, 208:47–57, 2014. ISSN 0001-8686. doi? https://doi.org/10.1016/j.cis.2014.02.002. URL http://www.sciencedirect.com/science/article/pii/S0001868614000360. Special issue in honour of Wolfgang Helfrich.

[18] Hammad A. Faizi, Shelli L. Frey, Jan Steinkühler, Rumiana Dimova, and Petia M. Vlahovska. Bending rigidity of charged lipid bilayer membranes. Soft Matter, 15:6006–6013, 2019. doi:10.1039/C9SM00772E. URL http://dx.doi.org/10.1039/C9SM00772E.

[19] Hélène Bouvrais, Flemming Cornelius, John H. Ipsen, and Ole G. Mouritsen. Intrinsic reaction-cycle time scale of na+,k+-atpase manifests itself in the lipid-protein interactions of nonequilibrium membranes. Proceedings of the National Academy of Sciences, 109(45):18442–18446, 2012. ISSN 0027-8424. doi:10.1073/pnas.1209909109. URL http://www.pnas.org/content/109/45/18442.

[20] Yuval Elani, Sowmya Purushothaman, Paula J. Booth, John M. Seddon, Nicholas J. Brooks, Robert V. Law, and Oscar Ces. Measurements of the effect of membrane asymmetry on the mechanical properties of lipid bilayers. Chem. Commun., 51:6976–6979, 2015. doi: 10.1039/C5CC00712G. URL http://dx.doi.org/10.1039/C5CC00712G.

[21] Víctor G. Almendro-Vedia, Paolo Natale, Michael Mell, Stephanie Bonneau, Francisco Monroy, Frederic Joubert, and Iván López-Montero. Nonequilibrium fluctuations of lipid membranes by the rotating motor protein f1f0-atp synthase. Proceedings of the National Academy of Sciences, 114(43):11291–11296, 2017. ISSN 0027-8424. doi:10.1073/pnas.1701207114. URL http://www.pnas.org/content/114/43/11291.

[22] P. Meleard, T. Pott, H. Bouvrais, and J. H. Ipsen. Advantages of statistical analysis of giant vesicle flickering for bending elasticity measurements. EUROPEAN PHYSICAL JOURNAL E, 34(10), OCT 2011. ISSN 1292-8941. doi:10.1140/epje/i2011-11116-6.

[23] Dominik Drabik, Magda Przybyłlo, Grzegorz Chodaczek, Ales Iglič, and Marek Langner. The modified fluorescence based vesicle fluctuation spectroscopy technique for determination of lipid bilayer bending properties. Biochimica et Biophysica Acta (BBA) - Biomembranes, 1858(2):244–252, 2016. ISSN 0005-2736. doi: https://doi.org/10.1016/j.bbamem.2015.11.020. URL http://www.sciencedirect.com/science/article/pii/S0005273615003922.

[24] Joanna B. Dahl, Vivek Narsimhan, Bernardo Gouveia, Sanjay Kumar, Eric S. G. Shaqfeh, and Susan J. Muller. Experimental observation of the asymmetric instability of intermediate-reduced-volume vesicles in extensional flow. Soft Matter, 12:3787–3796, 2016. doi: 10.1039/C5SM03004H. URL http://dx.doi.org/10.1039/C5SM03004H.

[25] S. Alex Rautu, Davide Orsi, Lorenzo Di Michele, George Rowlands, Pietro Cicuta, and Matthew S. Turner. The role of optical projection in the analysis of membrane fluctuations. Soft Matter, 13:3480–3483, 2017. doi: 10.1039/C7SM00108H. URL http://dx.doi.org/10.1039/C7SM00108H.

[26] Andrew F. Loftus, Sigrid Noreng, Vivian L. Hsieh, and Raghuveer Parthasarathy. Robust measurement of membrane bending moduli using light sheet fluorescence imaging of vesicle fluctuations. Langmuir, 29(47): 14588–14594, 2013. doi:10.1021/la403837d. URL https://doi.org/10.1021/la403837d. PMID: 24180269.

[27] V. Vitkova, J. Genova, M. D. Mitov, and I. Bivas. Sugars in the aqueous phase change the mechanical properties of lipid mono- and bilayers. Molecular Crystals and Liquid Crystals, 449(1):95–106, 2006. doi: 10.1080/15421400600582515. URL https://doi.org/10.1080/15421400600582515.

[28] Marzieh Karimi, Jan Steinkuhler, Debjit Roy, Raktim Dasgupta, Reinhard Lipowsky, and Rumiana Dimova. Asymmetric ionic conditions generate large membrane curvatures. Nano Letters, 18(12):7816–7821, 2018. doi: 10.1021/acs.nanolett.8b03584. URL https://doi.org/10.1021/acs.nanolett.8b03584. PMID: 30456959.

[29] J. R. Henriksen and J. H. Ipsen. Thermal undulations of quasi-spherical vesicles stabilized by gravity. European Physical Journal E, 9(4):365–374, 2002. ISSN 1292-8941. URL <GotoISI>://000181390400008.

[30] Hélène Bouvrais, Tanja Pott, Luis A. Bagatolli, John H. Ipsen, and Philippe Méléard. Impact of membrane-anchored fluorescent probes on the mechanical properties of lipid bilayers. Biochimica et Biophysica Acta (BBA) - Biomembranes, 1798 (7):1333–1337, 2010. ISSN 0005-2736. doi: https://doi.org/10.1016/j.bbamem.2010.03.026. URL http://www.sciencedirect.com/science/article/pii/S0005273610001252. Microscopy Imaging of Membrane Domains.

[31] Hélène Bouvrais, Lars Duelund, and John H. Ipsen. Buffers affect the bending rigidity of model lipid membranes. Langmuir, 30(1):13–16, Jan 2014. ISSN 0743-7463. doi:10.1021/la403565f. URL https://doi.org/10.1021/la403565f.

[32] Aidan T. Brown, Jurij Kotar, and Pietro Cicuta. Active rheology of phospholipid vesicles. Phys. Rev. E, 84:021930, Aug 2011. doi:10.1103/PhysRevE.84.021930. URL https://link.aps.org/doi/10.1103/PhysRevE.84.021930.

[33] U. Seifert. Fluid membranes in hydrodynamic flow fields: Formalism and an application to fluctuating quasispherical vesicles. Eur. Phys. J. B, 8:405–415, 1999.

[34] L. Miao, M. A. Lomholt, and J. Kleis. Dynamics of shape fluctuations of quasi-spherical vesicles revisited. Eur. Phys. J. E., 9:143–160, 2002.

[35] H. Zhou, B. B. Gabilondo, W. Losert, and W. van de Wate. Stretching and relaxation of vesicles. Phys. Rev. E, 83:011905, 2011.

[36] R. Lipowsky. Chapter 11 - generic interactions of flexible membranes. In R. Lipowsky and E. Sackmann, editors, Structure and Dynamics of Membranes, volume 1 of Handbook of Biological Physics, pages 521–602. North-Holland, 1995. doi:https://doi.org/10.1016/S1383-8121(06)80004-7. URL http://www.sciencedirect.com/science/article/pii/S1383812106800047.

[37] R. Dimova. Recent developments in the field of bending rigidity measurements on membranes. Adv. Coll. Int. Sci., 208:225–234, 2014.

[38] R. S. Gracia, N. Bezlyepkina, R. L. Knorr, R. L. Lipowsky, and R. Dimova. Effect of cholesterol on the rigidity of saturated and unsaturated membranes: fluctuation and electrodeformation analysis of giant vesicles. Soft Matter, 6:1472–1482, 2010.

[39] Dinesh Kumar, Channing M. Richter, and Charles M. Schroeder. Conformational dynamics and phase behavior of lipid vesicles in a precisely controlled extensional flow. Soft Matter, 16:337–347, 2020. doi: 10.1039/C9SM02048A. URL http://dx.doi.org/10.1039/C9SM02048A.

[40] Aidan T. Brown, Jurij Kotar, and Pietro Cicuta. Active rheology of phospholipid vesicles. Phys. Rev. E, 84:021930, Aug 2011. doi:10.1103/PhysRevE.84.021930. URL https://link.aps.org/doi/10.1103/PhysRevE.84.021930.

[41] Arwen I. I. Tyler, Jake L. Greenfield, John M. Seddon, Nicholas J. Brooks, and Sowmya Purushothaman. Coupling phase behavior of fatty acid containing membranes to membrane bio-mechanics. Frontiers in Cell and Developmental Biology, 7:187, 2019. ISSN 2296-634X. doi:10.3389/fcell.2019.00187. URL https://www.frontiersin.org/article/10.3389/fcell.2019.00187.

[42] Sowmya Purushothaman, Pietro Cicuta, Oscar Ces, and Nicholas J. Brooks. Influence of high pressure on the bending rigidity of model membranes. The Journal of Physical Chemistry B, 119(30):9805–9810, Jul 2015. ISSN 1520-6106. doi:10.1021/acs.jpcb.5b05272. URL https://doi.org/10.1021/acs.jpcb.5b05272.

[43] P Shchelokovskyy, S Tristram-Nagle, and R Dimova. Effect of the HIV-1 fusion peptide on the mechanical properties and leaflet coupling of lipid bilayers. New Journal of Physics, 13(2):025004, feb 2011. doi:10.1088/1367-2630/13/2/025004. URL https://doi.org/10.1088%2F1367-2630%2F13%2F2%2F025004.

[44] S. Alex Rautu, Davide Orsi, Lorenzo Di Michele, George Rowlands, Pietro Cicuta, and Matthew S. Turner. The role of optical projection in the analysis of membrane fluctuations. Soft Matter, 13:3480–3483, 2017. doi: 10.1039/C7SM00108H. URL http://dx.doi.org/10.1039/C7SM00108H.

[45] W. Rawicz, K.C. Olbrich, T. McIntosh, D. Needham, and E. Evans. Effect of chain length and unsaturation on elasticity of lipid bilayers. Biophys. J., 79:328–339, 2000.

[46] Jianjun Pan, Stephanie Tristram-Nagle, Norbert Kučerka, and John F. Nagle. Temperature dependence of structure, bending rigidity, and bilayer interactions of dioleoylphosphatidylcholine bilayers. Biophysical Journal, 94(1):117–124, Jan 2008. ISSN 0006-3495. URL http://www.sciencedirect.com/science/article/pii/S0006349508707707.

[47] M. S. Jablin, K. Akabori, and J. F. Nagle. Experimental support for tilt-dependent theory of biomembrane mechanics. Phys. Rev. Lett., 113:248102, Dec 2014. doi: 10.1103/PhysRevLett.113.248102. URL https://link.aps.org/doi/10.1103/PhysRevLett.113.248102.

[48] Sudipta Gupta, Judith U. De Mel, Rasangi M. Perera, Piotr Zolnierczuk, Markus Bleuel, Antonio Faraone, and Gerald J. Schneider. Dynamics of phospholipid membranes beyond thermal undulations. The Journal of Physical Chemistry Letters, 9(11):2956–2960, Jun 2018. doi:10.1021/acs.jpclett.8b01008. URL https://doi.org/10.1021/acs.jpclett.8b01008.

[49] Judith U. De Mel, Sudipta Gupta, Rasangi M. Perera, Ly Ngo, Piotr Zolnierczuk, Markus Bleuel, Sai Venkatesh Pingali, and Gerald J. Schneider. Influence of salt on membrane rigidity of neu-tral dopc vesicles, 2020.

[50] T. Betz and C. Sykes. Time resolved membrane fluctuation spectroscopy. Soft Matter, 8:5317–5326, 2012.

[51] Miglena I. Angelova and Dimiter S. Dimitrov. Liposome electroformation. Faraday Discuss. Chem. Soc., 81:303–311, 1986. doi:10.1039/DC9868100303. URL http://dx.doi.org/10.1039/DC9868100303.

[52] Ken Kelley Scott E. Maxwell, Harold D. Delaney. Designing Experiments and Analyzing Data: A Model Comparison Perspective. Taylor and Francis, 3 edition, 2017.

[53] Michael Chernick. Bootstrap Methods: A Guide for Practitioners and Researchers. John Wiley & Sons, Inc., 2 edition, 2007.

[54] U. Seifert. Configurations of fluid membranes and vesicles. Advances in physics, 46:13–137, 1997.

[55] S. T. Milner and S. A. Safran. Dynamical fluctuations of droplet microemulsions and vesicles. Phys. Rev. A, 36: 4371–4379, 1987.

[56] P. M. Vlahovska. Dynamics of membrane bound particles: capsules and vesicles. In C. Duprat and H.A. Stone, editors, Low-Reynolds-Number Flows: Fluid-Structure Interactions. Royal Society of Chemistry Series RSC Soft Matter, 2016.

[57] Petia M. Vlahovska and C. Misbah. Theory of vesicle dynamics in flow and electric fields. In R. Dimova and C. Marques, editors, The Giant Vesicle Book, page Chapter 7. CRC Press, 2019.

